# Modulation of Human T-type Calcium Channels by Synthetic Cannabinoid Receptor Agonists *in vitro*

**DOI:** 10.1101/2020.07.22.215434

**Authors:** Chris Bladen, Somayeh Mirlohi, Marina Santiago, Mitchell Longworth, Michael Kassiou, Sam Banister, Mark Connor

## Abstract

**BACKGROUND AND PURPOSE:** Consumption of Synthetic Cannabinoid Receptor agonists (SCRAs) is associated with severe adverse reactions including seizures, arrhythmias and death, but the molecular mechanisms surrounding SCRA toxicity are not yet established. These disease-like symptoms are also synonymous with altered T-type calcium channel activity which controls rhythmicity in the heart and brain. This study examined whether SCRAs alter T-type activity and whether this represents a possible mechanism of toxicity.

**EXPERIMENTAL APPROACH:** Fluorescence-based and electrophysiology assays were used to screen 16 structurally related synthetic cannabinoids for their ability to inhibit human T-type calcium channels expressed in HEK293 cells. The most potent compounds were then further examined using patch clamp electrophysiology.

**KEY RESULTS:** MDMB-CHMICA and AMB-CHMINACA potently blocked Cav3.2 with IC50 values of 1.5 and 0.74 μM respectively. Current inhibition increased from 47 to 80% and 45 to 87% respectively when the channel was in slow-inactivated state. Both SCRAs had little effect on steady state inactivation, however MDMB-CHMICA significantly shifted the half activation potential by −7mV. Neither drug produced frequency dependent block, in contrast to the phytocannabinoid Δ9-THC.

**CONCLUSIONS and IMPLICATIONS:** SCRAs are potent agonists of CB1 receptors and can be extremely toxic, but observed toxicity also resembles symptoms associated with altered Cav3.2 activity. Many SCRAs tested were potent modulators of Cav3.2, raising the possibility that SC toxicity may be due in part to Cav3.2 modulation. This potent T-type channel modulation suggests the possibility of SCRAs as a new drug class with potential to treat diseases associated with altered T-type channel activity.

## Introduction

The human genome includes three types of low-voltage-activated calcium channels (T-type *I*_ca_); Ca_v_3.1, Ca_v_3.2 and Ca_v_3.3 and each have specific cellular functions. (Perez-Reyes 2003). One of their key properties is their ability to activate near the threshold of cellular resting membrane potential, allowing quick responses to subtle changes in electrical activity and to amplify depolarizations, but also to inhibit them via afterhyperpolarizations (AHPs) following Ca^2^ influx (Bender *et al*., 2012). In particular, the Ca_v_3.2 channel is thought to play a critical role in many physiological processes by using these characteristics to control and regulate rhythmicity in the heart and brain (Kampa, Letzkus et al. 2007, David, Garcia et al. 2010, Ono and Iijima 2010, Bender, Uebele et al. 2012). Conversely, several disease states including cardiac arrhythmia, epilepsy and pain (Khosravani, Bladen et al. 2005, Heron, Khosravani et al. 2007, Ono and Iijima 2010, Zamponi, Lory et al. 2010) have been attributed to aberrant Ca_v_3.2 activity. Indeed in pain, up-regulation of Ca_v_3.2 has been linked to hyper-excitability of primary afferent fibers in chronic pain disorders, while animals with Ca_v_3.2 knocked out have shown dramatically reduced nociception in formalin assays of acute and inflammatory pain (Bourinet, Alloui et al. 2005, Choi, Na et al. 2007, Jagodic, Pathirathna et al. 2008, Bladen, McDaniel et al. 2015).

Synthetic Cannabinoid Receptor agonists (SCRAs) are a large class of drugs whose unregulated use and consumption in part aims to mimic the effects of Δ^9^-tetrahydrocannabinol (Δ^9^-THC), the principal psychoactive component of *Cannabis* (Banister and Connor 2018). However, some SCRAs are at least 300 times more efficacious at activating human cannabinoid (CB) receptor 1 when compared to Δ^9^-THC (Banister, Moir et al. 2015, Banister, Longworth et al. 2016, Sachdev, Vemuri et al. 2019). Not surprisingly, these highly potent drugs have been associated with many instances of hospitalization and even a significant number of deaths attributed to people consuming them (Banister, Longworth et al. 2016, Najafi, Dunn et al. 2016, Longworth, Connor et al. 2017). Although the adverse psychotomimetic effects seen in patients ingesting SCRAs are likely due to their interactions with CB receptors, it is less clear what causes the other severe adverse reactions such as abnormal heart rate, hyperalgesia and seizure activity (Trecki, Gerona et al. 2015, Najafi, Dunn et al. 2016, Tait, Caldicott et al. 2016, Babi, Robinson et al. 2017, EMCDDA 2017). In the library of SCRAs we tested for T-type channel activity in this report, two of the SC compounds MDMB-CHMICA and AMB-CHMINACA, potently modulated T-type calcium channel conductance (T-type *I*_ca_) in both a tonic and state-dependent manner. These two SCRAs have been identified as causing acute toxicity and death, yet they are not the most potent or efficacious activators of CB1 (Adamowicz 2016, Banister, Longworth et al. 2016, Najafi, Dunn et al. 2016, Sachdev, Vemuri et al. 2019).

To date, little is known about synthetic cannabinoid mechanisms of action beyond them being highly potent activators of cannabinoid receptors. This report has established that some synthetic cannabinoids also potently regulate and inhibit Ca_v_3.2, a calcium channel crucial to regulating heart and brain function. This modulation of a protein not traditionally thought of as a cannabinoid receptor not only adds to our basic understanding of synthetic cannabinoid interactions with human proteins, but it may point to an additional mechanism through which synthetic cannabinoids can cause acute toxicity in users.

## Methods

### Transfection and Cell culture

All experiments were performed using HEK293 FLPIN T-REX cells stably transfected with pcDNA5/FRT/TO constructs encoding human Ca_v_3.x cDNA as per Thermofisher protocol, (synthesized by Genscript,Piscataway, NJ, USA) together with pOG44 (Flp recombinase) plasmid using Fugene HD as per Promega protocol (Promega, Alexandria, NSW, Australia, Knapman *et al*., 2014a). Cells expressing the Ca_v_3.x constructs were selected using 150 μg/mL hygromycin + 15μg /mL blasticidin and cultured in DMEM supplemented with 10% FBS, 100 units/mL penicillin, 100 μg/mL streptomycin up until passage 5. Hygromycin was then reduced to 80 μg for subsequent passages. Ca_v_3.x expression was induced 24 h before FLIPR assays or electrophysiology experiments by adding 2μg/mL tetracycline. Cells were maintained and passaged at 80% confluency in 75-cm^2^ flasks and kept at 37°C/5% CO_2_. Cells for FLIPR assays were grown in same conditions but used at > 90% confluence.

### Electrophysiology

All whole-cell voltage-clamp recordings from HEK293 FLPIN T-REX cells stably transfected with human Ca_v_3.x were performed at room temperature. At least 24 h prior to experiments, cells were detached from flasks using trypsin/EDTA and plated into 10 cm sterile tissue culture dishes containing 10 mL of supplemented DMEM and 10-15 glass coverslips (12 mm diameter, ProScitech, QSLD, Australia). Culture dishes were then kept overnight in same conditions as flasks to allow cells to adhere to coverslips. They were then transferred to a 30°C/5% CO_2_ incubator to inhibit cell proliferation until ready to be used for electrophysiology experiments.

### Recording solutions

External recording solutions contained (in mM): 114 CsCl, 5 BaCl_2_, 1 MgCl_2_, 10 HEPES, 10 glucose, adjusted to pH 7.4 with CsOH. The internal patch pipette solution contained (in mM): 126.5 CsMeSO4, 2 MgCl_2_, 11 EGTA, 10 HEPES adjusted to pH 7.3 with CsOH. Internal solution was supplemented with 0.6 mM GTP and 2 mM ATP and mixed thoroughly just prior to use. Liquid junction potentials for above solutions were calculated prior to experiments using pClamp 10 software and corrected for during experiments. Compounds were prepared daily from 30 mM DMSO stocks and diluted into external solution just prior to use. Compounds were then applied rapidly and locally to the cells using a custom-built gravity driven micro-perfusion system (Feng, Doering et al. 2003). Initial vehicle experiments were performed to ensure that 0.1% DMSO had no effect on current amplitudes or channel kinetics (data not shown) and all subsequent experiments contained 0.1% DMSO in control external solutions. Currents were elicited from a holding potential of −100 mV and were measured by conventional whole-cell patch clamp techniques using an Axopatch 200B amplifier in combination with Clampex 9.2 software. (Molecular Devices, Sunnyvale, CA). After establishing whole cell configuration, cellular capacitance was minimized using the amplifier’s built-in analog compensation. Series resistance was kept to <10 MΩ and was compensated to at least 85% in all experiments. All data were digitized at 10 kHz with a Digidata 1320 interface (Molecular Devices) and filtered at 1 kHz (8-pole Bessel filter). Raw and online leak-subtracted data were both collected simultaneously, P/N4 leak subtraction was performed using opposite polarity and after the protocol sweep. For tonic inhibition of T-type current, membrane potential was stepped from −100 mV to −30 mV for 200 ms and then allowed to recover for 12 seconds (1 sweep). A minimum of 10 sweeps were collected under control external perfusion to allow for control peak current to equilibrate. Drug was then continuously perfused, and sweeps recorded until no further inhibition is seen (minimum of 3 sweeps with same amplitude). In current-voltage relation studies, the membrane potential was held at −100 mV and cells were depolarized from −70 to 50 mV in 10 mV increments. For steady-state inactivation studies, a 3.6 second conditioning pre-pulse of various magnitude (initial holding at −110 mV), was followed by a depolarizing pulse to −30 mV. Individual sweeps were separated by 12 seconds to permit recovery from inactivation between conditioning pulses. The duration of the test pulse was typically 20 ms and the current amplitude obtained from each test pulse was normalized to that observed at the holding potential of −100 mV.

### Data Analysis and Statistics

Data were acquired and analyzed using Clampfit 10.4 (Molecular Devices) and Origin 8.5 software (Northampton, MA, USA) or GraphPad Prism (Version 7.0b, San Diego, CA) was used in the preparation of all figures and curve fittings. Current-voltage (iV) relationships were fitted with a modified Boltzmann equation: I = [Gmax*(Vm-Erev)]/[1+exp((V0.5act-Vm)/ka)], where Vm is the test potential, V0.5act is the half-activation potential, Erev is the reversal potential and Gmax is the maximum slope conductance. Steady-state inactivation curves were fitted using Boltzmann equation: I =1/(1 + exp((Vm - Vh)/k)), where Vh is the half-inactivation potential and k is the slope factor. Dose-response curves were fitted with the equation y = A2 + (A1-A2)/(1 + ([C]/IC50)^n^) where A1 is initial current amplitude and A2 is the current amplitude at saturating drug concentrations, [C] is the drug concentration and n is the Hill coefficient. Statistical significance was determined using Student’s t-tests and one-way or repeated measures ANOVA. Significant values are indicated in text and figure legends. All data are given as means +/**-** standard errors. Correlation between FLIPR and Electrophysiology (Ephys) assays was determined as poor when values exceeded >50% difference between the two assays. When tonic inhibition of ion channel was greater than 80% in both assays, compounds were considered to be potent inhibitors of the calcium current and chosen for further testing of kinetic interactions using electrophysiological techniques.

### Assay of intracellular calcium concentration (FLIPR assay)

Changes in intracellular calcium ([Ca]i) were measured using a fluorometric imaging plate reader (FLIPR) and using Calcium 5 membrane potential assay kit (Molecular Devices Sunnyvale, CA). Briefly, 24 h prior to the assay, HEK-3.x cells were detached from the flask using trypsin/EDTA (Sigma-Aldrich) and resuspended in 10 mL Leibovitz’s L-15 media supplemented with 1% FBS, 100 U penicillin and 100 μg streptomycin per mL, 15 mM glucose 2 μg/mL tetracycline. Cells were plated in a volume of 90 μL per well in black-walled, clear-bottomed 96-well microplates (Corning, Castle Hill, Australia), and incubated overnight at 37°C in ambient CO_2_. Dye was reconstituted with assay buffer (HBSS) containing in mM: NaCl 145, HEPES 22, Na2HPO4 0.338, NaHCO3 4.17, KH2PO4 0.441, MgSO4 0.407, MgCl_2_ 0.493, CaCl_2_ 1.26, glucose 5.56, pH 7.4, osmolarity 315 mOSM. Cells were loaded with 90 μL per well of the dye solution without removal of the L-15 and incubated at 37°C for 60 minutes in ambient CO_2_. Fluorescence was measured using a FlexStation 3 microplate reader (Molecular Devices, excitation 530 nm/565 nm). Baseline readings were taken every 2s for at least 2 min, at which time drug was added in a volume of 20 μL to cells and fluorescence recorded for 5 min in drug solution. 10 mM CaCl_2_ was then added to activate the calcium channels and the fluorescence was recorded for a further 3 min. Area under the curve was measured and expressed as change in fluorescence elicited by addition of drug versus percentage of peak baseline fluorescence after subtraction of the changes produced by vehicle addition as previously described (Knapman, Santiago et al. 2013). Final concentration of vehicle (DMSO) was not more than 0.1%. Data were expressed as the mean ± SEM of at least 5 independent determinations performed in duplicate, unless otherwise stated. Pooled data were fit to a 4-parameter logistic equation in Graphpad PRISM 7 (GraphPad Software, San Diego CA).

### Drugs and reagents

Stock Drugs were dissolved in DMSO and diluted fresh before each use in external recording solution to give a final vehicle concentration of 0.1%. Synthetic Cannabinoids were synthesised using methods previously described (Parry, Eiblmaier et al. 2007). Cell culture media, buffers, antibiotics, and general chemicals were from Life Technologies, Sigma-Aldrich or InvivoGen (San Diego, CA, USA).

## Results

### Initial FLIPR screening of inhibition of T-type currents by Synthetic Cannabinoids

Inhibition of T-type calcium channels was measured using a Fluorescence Imaging Plate Reader (FLIPR) assay and results expressed as percentage of baseline fluorescence before and after addition of 10 μM synthetic cannabinoid receptor agonist. 8 SCRAs showed > 80% inhibition of the increase in fluorescence produced by addition of 10 mM Ca^2^ in cells expressing Ca_v_3.1. Similarly, 3 drugs produced >80% fluorescence inhibition of signal in cells expressing Ca_v_3.2 and Ca_v_3.3 (Table 1), but only AMB-CHMINACA produced >80% fluorescence inhibition of signal across all three T-type channels using the FLIPR assay (Table 1 and Figure 1B).

**Table 1.** Initial screening of Synthetic Cannabinoids using Fluorescence Imaging Plate Reader (FLIPR) and Patch Clamp Electrophysiology (Ephys). Table 1 shows results of 16 synthetic cannabinoids screened at 10 μM with both FLIPR and Ephys assays on stably expressed human T-type calcium channels in HEK293T Cells (hCa_v_ 3.1, hCa_v_ 3.2 and hCa_v_ 3.3). Compounds that produced greater than 80% fluorescence (FLIPR) or current inhibition (Ephys) at this concentration were designated as potent and highlighted in green. FLIPR identified 8 SCRAs that inhibited fluorescence >80% in hCa_v_ 3.,1 however Ephys produced only 1, MDMB-CHMINACA. In hCa_v_ 3.2, FLIPR identified three compounds MDMB-CHMICA, MDMB-FUBICA and AMB-CHMINACA as potent fluorescence inhibitors in hCa_v_ 3.2 while Ephys confirmed two of those compounds, MDMB-CHMICA and AMB-CHMINACA as potent current blockers. In hCa_v_ 3.3, FLIPR identified 3 SCRAs that showed >80% fluorescence inhibition, however in the Ephys assay, none of the SC compounds showed any significant block. Variance of greater than 50% between the FLIPR and Ephys assays are highlighted in red and Ephys data for phytocannabinoids’ **THC** and **CBD** are shown for reference purposes. Data are % fluorescence or current inhibition and represent the mean ± SEM of 6 independent FLIPR assays per compound on each stably expressed ion channel or the mean ± SEM of 6 independent whole-cell patch clamp recordings per compound on each channel.

|  | Structural subunits |  |  | FLIPR | Ephys | FLIPR | Ephys | FLIPR | Ephys |
| --- | --- | --- | --- | --- | --- | --- | --- | --- | --- |
| Compound Name | Core | Alkyl chain | Amino acid | Ca<sub>v</sub> 3.1 | Ca<sub>v</sub> 3.1 | Ca<sub>v</sub> 3.2 | Ca<sub>v</sub> 3.2 | Ca<sub>v</sub> 3.3 | Ca<sub>v</sub> 3.3 |
| AMB-CHMICA | Indole | (Cyclohexyl) methyl | Valinate | 81.8 ± 2.8 | 27.3 ± 1.17 | 72.3 ± 1.2 | 56.3 ± 7.9 | 85.5 ± 1.4 | 21.5 ± 2.9 |
| MDMB CHMICA | Indole | (Cyclohexyl) methyl | t-Leucinate | 85.8 ± 1.1 | 78.0 ± 3.2 | 89.7 ± 1.0 | 89.9 ± 1.7 | 60.1 ± 5.7 | 36.0 ± 2.1 |
| AMB-CHMINACA | Indazole | (Cyclohexyl) methyl | Valinate | 91.3 ± 1.1 | 64.2 ± 5.0 | 83.3 ± 0.6 | 81.0 ± 3.7 | 94.1 ± 0.6 | 29.9 ± 1.5 |
| MDMB-CHMINACA | Indazole | (Cyclohexyl) methyl | t-Leucinate | 75.1 ± 5.4 | 80.8 ± 0.9 | 56.8 ± 3.3 | 61.3 ± 7.1 | 33.7 ± 6.9 | 41.3 ± 5.4 |
| AMB-FUBICA | Indole | 4-Fluorobenzyl | Valinate | 41.3 ± 3.6 | 46.4 ± 1.6 | 53.9 ± 0.7 | 47.6 ± 8.6 | 6.1 ± 1.1 | 10.6 ± 3.6 |
| MDMB-FUBICA | Indole | 4-Fluorobenzyl | t-Leucinate | 95.8 ± 0.8 | 77.7 ± 3.7 | 94.7 ± 0.5 | 32.8 ± 8.4 | 52.8 ± 5.2 | 30.8 ± 4.0 |
| AMB-FUBINACA | Indazole | 4-Fluorobenzyl | Valinate | 92.2 ± 2.3 | 61.8 ± 3.9 | 69.6 ± 0.5 | 54.9 ± 9 | 18.2 ± 1.3 | 2.6 ± 0.6 |
| MDMB FUBINACA | Indazole | 4-Fluorobenzyl | t-Leucinate | 38.0 ± 1.3 | 46.8 ± 3.9 | 68.2 ± 1.3 | 45.3 ± 5.7 | 4.8 ± 3.7 | 33.1 ± 5.4 |
| AMB-PICA | Indole | Pentyl | Valinate | 38.3 ± 2.5 | 28.0 ± 5.8 | 15.8 ± 5.6 | 35.1 ± 7.3 | 13.4 ± 2.4 | 13.3 ± 4.1 |
| MDMB PICA | Indole | Pentyl | t-Leucinate | 5.5 ± 1.7 | 13.6 ± 2.2 | 58.5 ± 2.9 | 40.3 ± 6.5 | 8.7 ± 2.0 | 6.7 ± 2.2 |
| AMB-PINACA | Indazole | Pentyl | Valinate | 95.3 ± 0.9 | 54.5 ± 2.6 | 83.7 ± 1.2 | 45.9 ± 3.4 | 95.5 ± 1.1 | 26.9 ± 2.0 |
| MDMB-PINACA | Indazole | Pentyl | t-Leucinate | 92.7 ± 2.4 | 67.8 ± 5.5 | 59.6 ± 3.3 | 46.1 ± 9.5 | 55.4 ± 7.8 | 38.0 ± 4.0 |
| 5F-AMB-PICA | Indole | 5-Fluoropentyl | Valinate | 14.4 ± 2.4 | 12.1 ± 1.9 | 13.5 ± 3.6 | 22.5 ± 3.5 | 10.8 ± 1.9 | 2.1± 0.8 |
| 5F-MDMB-PICA | Indole | 5-Fluoropentyl | t-Leucinate | 51.4 ±1.4 | 42.6 ± 2.1 | 66.3 ± 1.5 | 48.5 ± 0.8 | 11.9 ± 3.7 | 8.1 ± 2.1 |
| 5F-AMB-PINACA | Indazole | 5-Fluoropentyl | Valinate | 28.8 ± 6.0 | 27.6 ± 2.8 | 13.5 ± 3.6 | 44.1 ± 9.5 | 4.2 ± 4.2 | 2.2 ± 0.5 |
| 5F-MDMB-PINACA (5F-ADB) | Indazole | 5-Fluoropentyl | t-Leucinate | 81.4 ± 0.6 | 67.6 ± 1.6 | 56.8 ± 1.5 | 43.5 ± 1.8 | 24.0 ± 3.8 | 24.0 ± 3.5 |
| Δ<sup>9</sup>-tetrahydrocannabinol | - | - | - |  | 74.9 ± 3.9 |  | 43.4 ± 5.5 |  | 6.4 ± 3.1 |
| Cannabidiol | - | - | - |  | 69.5 ± 4.0 |  | 48.0 ± 5.2 |  | 31.6 ± 4.8 |

**Figure 1.**
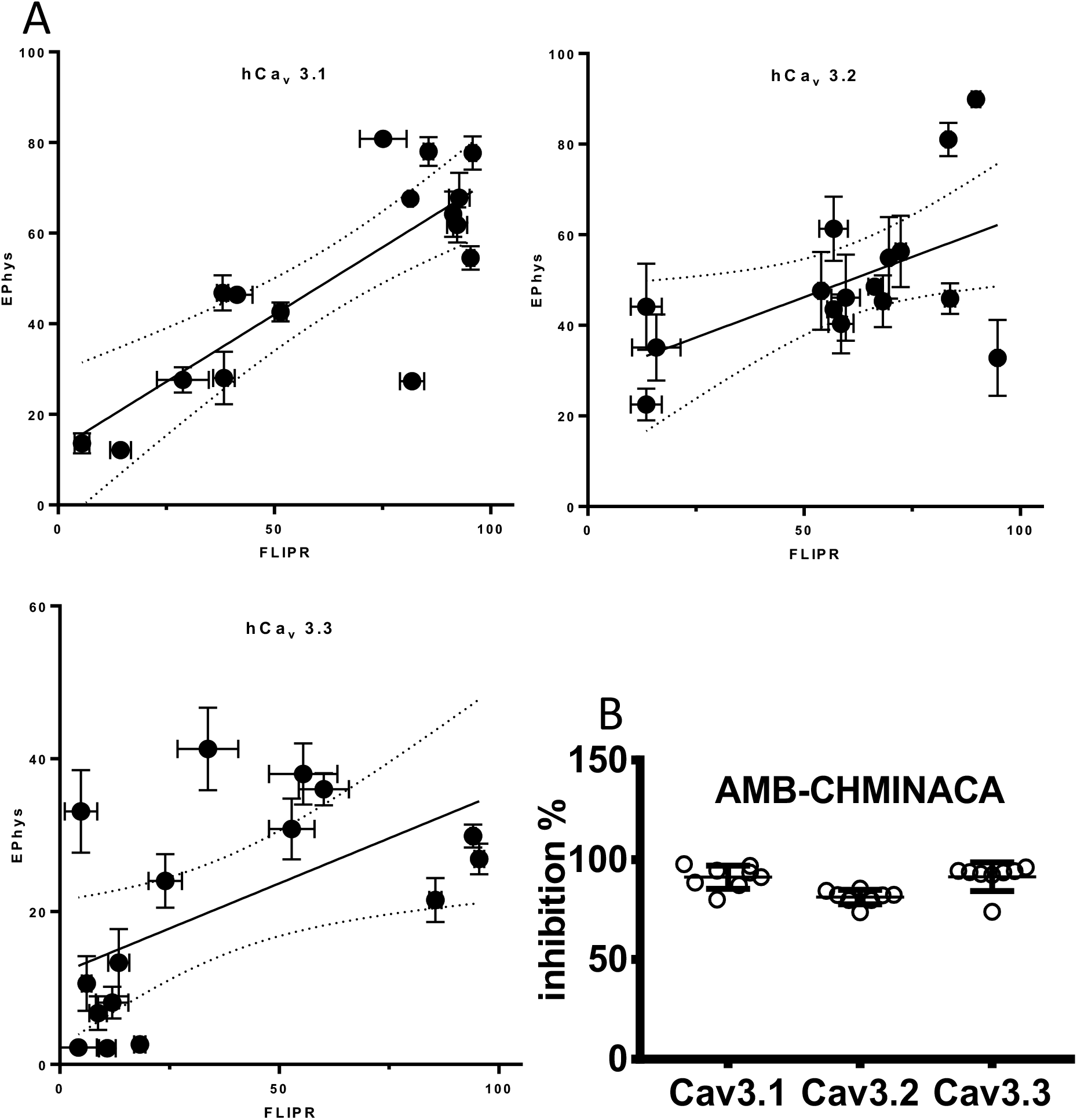
Comparison of Fluorescence Imaging Plate Reader (FLIPR) screening of Synthetic Cannabinoids to Patch Clamp Electrophysiology. 1A. Results of the FLIPR assay (X axis) were plotted against the results of the Ephys assay (Y axis) and a linear regression analysis performed between the 2 sets of data. Overall, the correlation between synthetic cannabinoid screening (10 μM) with FLIPR and Ephys assays on all 3 stably expressed human T-type calcium channels was excellent with R values of 0.66, 0.30 and 0.32 and P values of 0.0001, 0.030 and 0.023 for hCa_v_ 3.1, hCa_v_ 3.2 and hCa_v_ 3.3 respectively. Of the sixteen SCRAs tested, only AMB-CHMICA in hCa_v_ 3.1 and MDMB-FUBICA in hCa_v_ 3.2 had a variance of greater than 50% between the two assays (also highlighted red in Table 1). In addition, when screening hCa_v_ 3.3, the FLIPR assay identified 3 compounds as potent inhibitors, however none of the SCRAs showed any significant block in the Ephys assay. 1B. Results of the FLIPR data for AMB-CHMINACA are shown since this was the only compound to show potent % fluorescence inhibition across all three T-type calcium channels.

### Initial Electrophysiology screening of tonic inhibition of T-type currents by Synthetic Cannabinoids

Tonic inhibition of T-type currents was measured by stepping the cell membrane potential repetitively from −100 mV to −30 mV (sweep) to elicit peak current amplitude while applying continuous perfusion of 10 μM synthetic cannabinoid until inhibition reached steady state which was deemed to be obtained when no further reduction in current amplitude was observed after 3 successive sweeps (36 seconds). Again, a threshold of 80% inhibition of the initial current was used to identify a compound as having significant inhibitory effects on the Ca_v_3.x channels. Screening of the 16 SCRAs identified 3 compounds, MDMB-CHMICA, MDMB-CHMINACA and MDMB-FUBICA as potent inhibitors of Ca_v_3.1. For Ca_v_3.2, MDMB-CHMICA and AMB-CHMINACA were potent inhibitors, however none of the drugs produced an inhibition of Ca_v_3.3 > 80% in these conditions (Table 1).

Overall, the correlation between high throughput FLIPR and electrophysiology screening of synthetic cannabinoids was excellent (Figure 1A). Of the sixteen synthetic cannabinoid compounds screened using both methods, only AMB-CHMICA in Ca_v_3.1 and MDMB-FUBICA in Ca_v_3.2, did not correlate (Table 1 highlighted in red). In Ca_v_3.3, three compounds AMB-PINICA, AMB-CHMINACA and AMB-CHMICA produced significant inhibition of the fluorescence signal in the FLIPR assay, however the Ephys results showed minimal tonic inhibition of current by these 3 compounds (Table 1 highlighted in red). It is important to note that the results of the other 13 synthetic cannabinoids in Ca_v_3.3 showed excellent correlation between the two assays.

Finally, we also tested the two best-known phytocannabinoids, CBD and Δ^9^-THC, using electrophysiology (Table 1) and they inhibited T-type *I*_ca_ with results in excellent agreement with previous findings (Ross, Napier et al. 2008).

Following on from the preliminary screening, we further examined the effects of the 2 most potent inhibitors of Ca_v_3.2, MDMB-CHMICA and AMB-CHMINACA on the biophysical properties of these channels *in vitro*. Using a high concentration (10 μM) of both MDMB-CHMICA and AMB-CHMINACA to measure time course of Ca_v_3.2 current inhibition, results showed both compounds rapidly inhibited Ca_v_3.2 current amplitude. However, during the washout phase, MDMB-CHMICA inhibition was almost completely reversed after 2 min, whereas AMB-CHMINACA achieved only <30% reversal of inhibition after 3 min (Figure 2).

**Figure 2.**
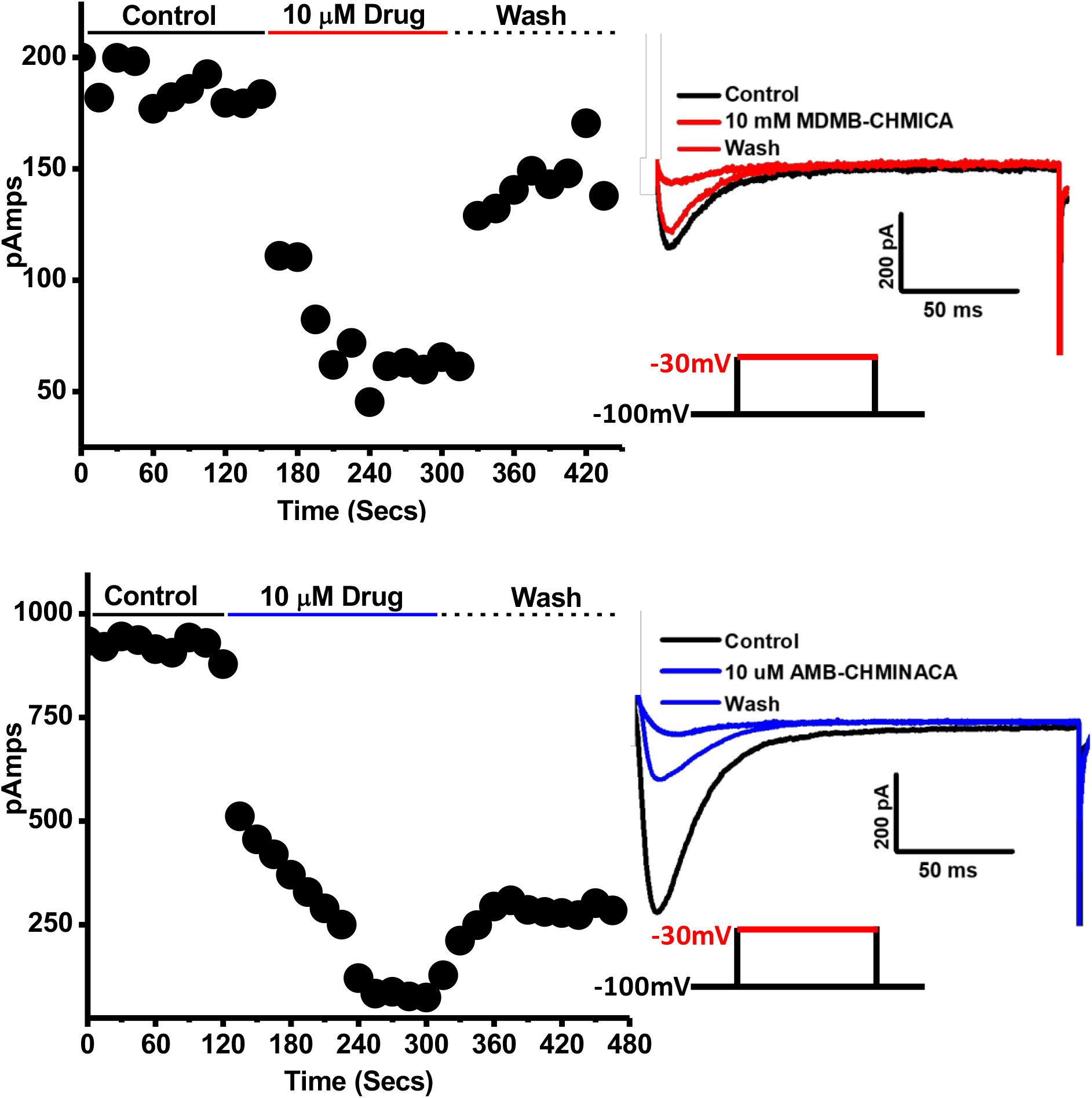
Representative time course of hCa_v_ 3.2 current inhibition by MDMB-CHMICA and AMB-CHMINACA. Using the same step voltage protocol (inset) and same drug concentration (10 μM) as the Ephys screening assay, time course of hCa_v_3.2 inhibition was recorded followed by washout. Both drugs rapidly blocked hCa_v_3.2, however MDMB-CHMICA (Red) block was almost completely reversed on washout whereas AMB-CHMINACA (Blue) washout was <30%. Right panels are current traces from the endpoint of control, drug and wash.

To calculate the half maximal inhibitory concentration (IC50) of both SCRAs on Ca_v_3.2 in tonic block conditions, concentration-response curves were generated by superfusing one concentration of drug onto an isolated cell and recording the percent inhibition compared to current measured under vehicle perfusion on the same cell. Data were pooled from at least six independent determinations for each drug concentration and fit to a logistic equation (see methods) which yielded IC50 values of 1.5 ± 0.2 μM and 0.74 ± 0.3 μM for MDMB-CHMICA & AMB-CHMINACA respectively (Figure 3). Using the same tonic inhibition protocol, Δ^9^-THC and CBD produced slightly less than 50% inhibition at 10 μM in the screening assay (Table 1), suggesting the two SCs may be up to 10-fold more potent at inhibiting Ca_v_3.2 conductance under these conditions.

**Figure 3.**
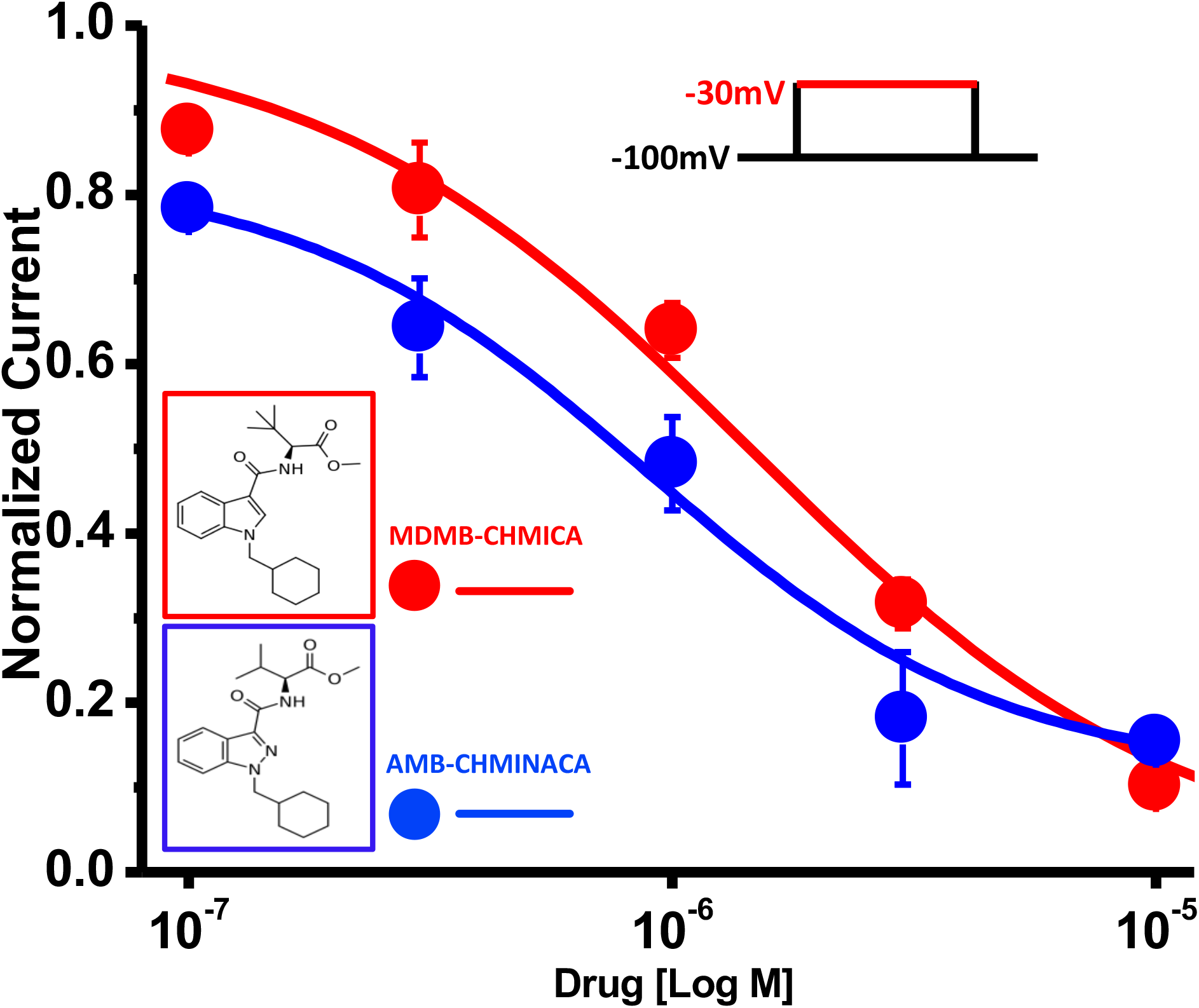
hCa_v_ 3.2 Dose Response Curves (IC50) of MDMB-CHMICA and AMB-CHMINACA. Dose-response curves were generated using Log concentrations of **MDMB-CHMICA (Red)** & **AMB-CHMINACA** (Blue) and fitted with Boltsmann iV equation to yield IC50 of 1.5 ± 0.2 & 0.74 ± 0.3 μM respectively (Current inhibition was normalised to vehicle control and represent the mean ± SEM of 6 independent cellular recordings)

### Effects of MDMB-CHMICA and AMB-CHMINACA on biophysical properties of Ca_v_3.2

The effects of MDMB-CHMICA and AMB-CHMINACA (1 μM each) on the voltage dependence of Ca_v_3.2 activation and inactivation were studied as described in the Methods. MDMB-CHMICA significantly shifted half-activation potential (V_0.5_) of Ca_v_3.2, (−54.8 ± 1 mV vs −47 ± 1 mV Control) (n=6 for all experiments). However, AMB-CHMINACA did not significantly alter V_0.5_ (−43 ± 1 mV vs −42.1 ± 1 mV control). Steady state inactivation was not affected by either MDMB-CHMICA (−78.6 ± 1 mV vs −77.1 ± 2 mV Control) or AMB-CHMINACA −77.1 ± 1 mV vs −80.2 ± 1 mV control) (Figures 4A and 4B). The effects of the synthetic cannabinoids on channel inactivation is in contrast with those of Δ^9^-THC and CBD, which have both been shown to significantly alter this parameter. However for activation kinetics, the significant negative shift by MDMB-CHMICA is similar to the negative shift caused by Δ^9^-THC, whereas AMB-CHMINACA is similar to CBD in that it does not significantly alter activation (Ross, Napier et al. 2008).

**Figure 4.**
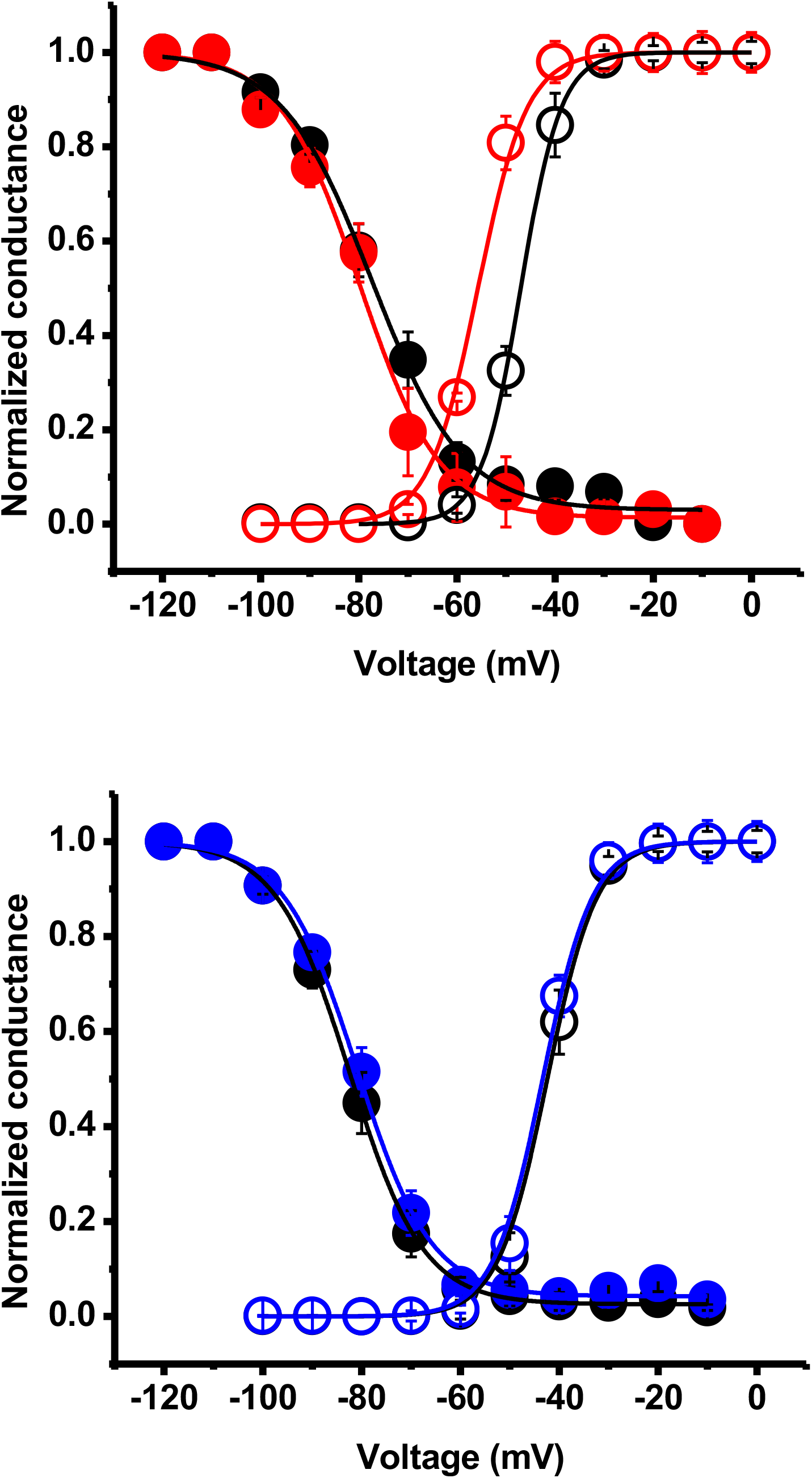
hCa_v_ 3.2 Activation and Inactivation kinetics of MDMB-CHMICA and AMB-CHMINACA. 4A. Using a current-voltage relationship protocol (iV), 1 μM **MDMB-CHMICA** significantly shifted half-activation potential **(v_0.5_)** of hCa_v_3.2 (−54.8 ± 1 mV vs −47 ± 1 mV Control) (Red open circles). However, Steady State Inactivation (SSI) was not affected (−78.6 ± 1 mV vs −77.1 ± 2 mV Control) (Red closed circles). 4B. Using the same protocols, 1 μM **AMB-CHMINACA** did not significantly alter **v_0.5_** (−43 ± 1 mV vs −42.1 ± 1 mV Control) (Blue open circles) or SSI (−77.1 ± 1 mV vs −80.2 ± 1 mV Control) (Blue closed circles). Results are normalised to peak current and represent the mean ± SEM of 6 independent cellular recordings).

Previously, it has been shown that compounds can preferentially block Ca_v_3.2 when the channels are in a slow-inactivated conformation (Bladen and Zamponi 2012). Therefore, block of Ca_v_3.2 by accumulating slow inactivation was assessed in the presence and absence of MDMB-CHMICA and AMB-CHMINACA (1 μM) using a 2-pulse voltage protocol where an initial depolarizing test pulse from −120 mV to −30 mV is used to evoke peak current (P1). This is followed by a 10 second pre-conditioning pulse to – 80 mV that removes the fast steady state inactivation, but maintains the channel in a prolonged slow inactivated state. This is then followed by another test pulse from −120 to −30 mV (P2). Any differences in inhibition between P1 and P2 peak current amplitude are considered to be attributed to drug preferentially binding while the channel is held in the slow-inactivated conformation. Using this protocol produced a significant increase in inhibition by both MDMB-CHMICA and AMB-CHMINACA with increases from 47% ±3.9 to 80% ±3.2 & 45% ±1.8 to 87% ±1.2 respectively, suggesting these drugs preferentially bind while in a slow inactivated state (Figure 5A). Similarly, Δ^9^-THC and CBD (10 μM) inhibition of Ca_v_3.2 was also significantly enhanced (from 43% ±2.4 to 71% ± 4 and 57% ±5 to 96% ±2 respectively), when channels were held in this conformation (Figure 5B).

**Figure 5.**
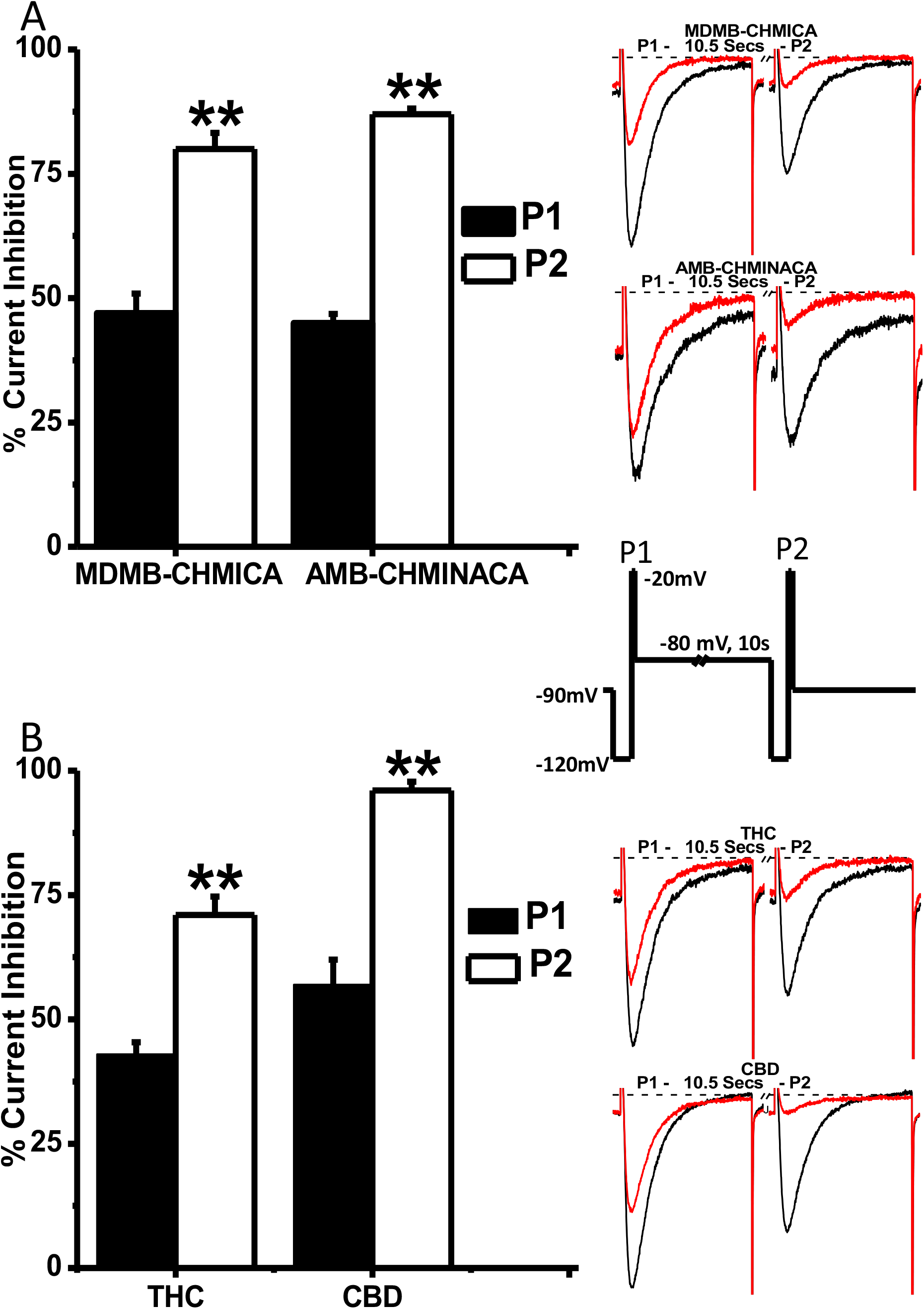
hCa_v_ 3.2 current inhibition in slow inactivated state by MDMB-CHMICA, AMB-CHMINACA, THC and CBD. 5A. A 2-pulse voltage protocol (Inset) was used to elicit hCa_v_3.2 into a slow inactivation state. This produced a dramatic increase in current inhibition by both **MDMB-CHMICA** & **AMB-CHMINACA** at 1 μM. For **MDMB-CHMICA**, current inhibition increased from 47 ± 3.9 % tonic (P1), to 80 ± 3.2 % in slow inactivated state (P2) and for **AMB-CHMINACA**, from 45 ± 1.8 to 87 ± 1.1 %. 5B. For THC, current inhibition increased from 43 ± 2.4 % tonic (P1), to 71 ± 3.7 % in slow inactivated state (P2) and for CBD, inhibition increased from 57 ± 5 % tonic (P1), to 96 ± 1.8 % in slow inactivated state (P2). Representative traces from P1 vs P2 for each drug are shown inset. Results are represented as percent inhibition of peak current of vehicle control and represent the mean ± SEM of 6 independent cellular recordings. Statistical analysis was performed using Student t-test with asterisks denoting significance at *p<0.05, **p<0.01, ***p<0.001

Many local anesthetics and anti-epileptic drugs currently used as therapies to target ion channels, exhibit preferential binding and efficacy in a frequency dependent manner (Sheets, Heers et al. 2008). We therefore examined frequency-dependent inhibition of Ca_v_3.2 by MDMB-CHMICA and AMB-CHMINACA at 0.5 and 1 Hz. However, when we evoked currents at 0.5 and 1 Hz, neither drug showed any significant change in the rate of inhibition vs vehicle control. (Figure 6A). In contrast, Δ^9^-THC showed a significant increase in inhibition at both 0.5 and 1 Hz vs vehicle control, whereas CBD showed no increase in inhibition (Figure 6B).

**Figure 6.**
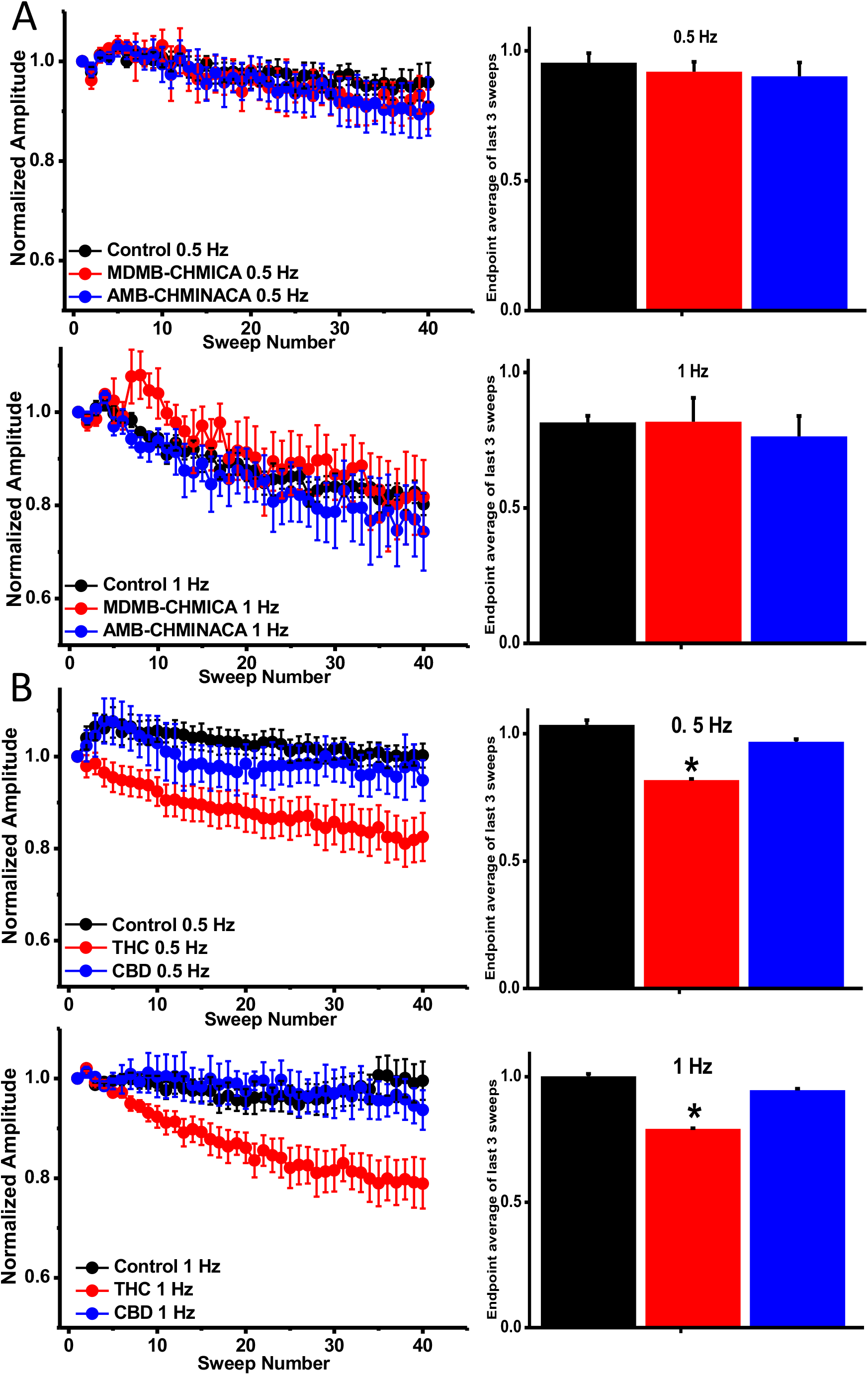
hC_av_ 3.2 Frequency dependent block of MDMB-CHMICA, AMB-CHMINACA, THC and CBD. 6A. Cells were held at −100 mV and stepped to-30 mV for 2 different frequencies (1 and 0.5 Hz), with inter-sweep holding potential of −100 mV. Peak currents from each sweep were normalized to the average peak current of first three sweeps and plotted against sweep number. Black symbols represent control vehicle without drugs, while red and blue symbols represent experiments in the presence of 1 μM **MDMB-CHMICA** and **AMB-CHMINACA**, respectively. Bar graphs represent the relative amplitudes of the average of the last 3 sweeps for each frequency. At 0.5 and 1 Hz (2 and 1 second inter-sweep intervals), neither drug showed significant difference from control. 6B. At 10 μM, THC but not CBD, showed a significant increase in frequency dependent inhibition after 40 sweeps. At 0.5 Hz and 1 Hz, inhibition increased 18.2 ± 0.4 % and 20.8 ± 0.3 % respectively. Statistical analysis was performed using Student t-test with asterisks denoting significance at *p<0.05.

## Discussion

The principal finding of this study is that synthetic cannabinoids MDMB-CHMICA and AMB-CHMINACA potently modulated T-type *I*_ca_ in both a tonic and state dependent manner. MDMB-CHMICA and AMB-CHMINACA effects on Ca_v_3.2 activity were distinct from each other, and quite different to the effects of the phytocannabinoids Δ^9^-THC and CBD. In particular, neither MDMB-CHMICA or AMB-CHMINACA altered the voltage-dependence of the inactivation of Ca_v_3.2, although they did potentiate the accumulation of slow channel inactivation, which can presumably contribute to the inhibition of channel current. The lack of effect on steady state channel inactivation is in contrast to the effects of Δ^9^-THC, CBD and endocannabinoids, which all produce profound hyperpolarizing shifts in the membrane potential at which Ca_v_3.2 opens (Chemin, Monteil et al. 2001, Ross, Napier et al. 2008, Ross, Gilmore et al. 2009, Gilmore, Heblinski et al. 2012). By contrast, MDMB-CHMICA, like Δ^9^-THC, shifted Ca_v_3.2 activation to more negative potentials, while AMB-CHMINACA, like CBD, had no such effect.

Several potential drug interaction sites have been identified on Ca_v_3 channels. A crystal structure of Ca_v_3.1 bound to the non-selective T-channel inhibitor Z944 (Zhao, Huang et al. 2019) showed the drug interacting with structural elements of the pore, as well as regions of the channel between domains II and III – so-called fenestrations, previously implicated in the inhibitory actions of local anaesthetic-like drugs at Ca_v_3 and Na_v_ channels (Bladen and Zamponi 2012, Martin and Corry 2014). The actions of Z944 are voltage-dependent, but it is not known whether the drug shifts the apparent voltage-dependence of steady state activation or inactivation of the channel, and its inhibitory effects were suggested to involve block of the channel pore by an aromatic moiety (Lee 2014). The block of Ca_v_3.2 in the absence of effects on steady state inactivation by both MDMB-CHMICA and AMB-CHMINACA suggest that they may also interact with the channel pore through their cyclohexyl moiety. The tert-leucine moiety of MDMB-CHMICA resembles the tert-butyl group of Z944, which interacts with the domain II-III fenestration in Ca_v_3.1 (Zhao, Huang et al. 2019). The only difference in structure between MDMB-CHMICA and AMB-CHMINACA is an extra methyl in the tert-leucine moiety of MDMB-CHMICA compared to the valine of AMB-CHMINACA. It is tempting to speculate that this slight difference may be responsible for the selective effects of AMB-CHMINACA on channel activation.

Both MDMB-CHMICA and AMB-CHMINACA were widely available unregulated drugs in Europe and the United States and have been associated with thousands of serious intoxications and dozens of deaths (EMCDDA 2017). Both drugs are high efficacy CB1 agonists relative to Δ^9^-THC (Sachdev, Vemuri et al. 2019), and *prima facie* the psychoactive effects of these drugs are likely due to this CB1 activity. CB1 agonist activity drives human use and almost certainly contributes to the toxicity of MDMB-CHMICA, AMB-CHMINACA and related drugs, but the role of CB1 activation in human intoxication idea remains untested. Intoxication by synthetic cannabinoids is associated with a myriad of symptoms, with many of the effects apparently occurring in the central nervous system. It is reasonable to attribute these to activation of CB1. In humans, the subjective psychological effects of Δ^9^-THC (Micale, Drago et al. 2019)) are significantly attenuated by the CB1 antagonists rimonabant and TM38837, which provides strong evidence for a contribution of CB1 receptors to the psychoactive responses to moderate doses of a low efficacy CB1 agonist and make it likely that synthetic cannabinoids exert CB1-dependent effects on mood. Similarly, the characteristic Δ^9^-THC-induced tachycardia observed in humans is blocked by rimonabant and TM38837, (Klumpers, Fridberg et al. 2013) (Micale, Drago et al. 2019) demonstrating a role for CB1 in alterations of some cardiovascular parameters. The concentration of synthetic cannabinoids detected in the serum of people who present for treatment varies tremendously, but for MDMB-CHMICA concentrations of up to 230 nM (86-91 ng/mL) have been reported in case series (Backberg, Tworek et al. 2017, Franz, Angerer et al. 2017), which at least raises the potential involvement of synthetic cannabinoid interactions with T-type calcium channels in toxicity. Intriguingly, a unique physiological consequence in humans of pharmacological inhibition of T-type calcium channels with the anti-hypertensive drug mibefradil was bradycardia, which has been reported to be associated with intoxication by MDMB-CHMICA and AMB-CHMINACA in a significant minority of cases (Hill, Najafi et al. 2016, Backberg, Tworek et al. 2017, Hermanns-Clausen, Muller et al. 2018). This is in contrast to the acute effects of Δ^9^-THC, and also many (but not all) other cases of MDMB-CHMICA and AMB-CHMINACA intoxication where tachycardia was a major presentation (Najafi, Dunn et al. 2016, Backberg, Tworek et al. 2017, Hermanns-Clausen, Muller et al. 2018). T-type Calcium channels and CB receptors are involved in many aspects of normal function, but the absence of studies in humans that differentiate the effects of modulating either protein in the brain makes it difficult to ascribe any aspects of SC toxicity to modulation of either.

In this study, our initial screening experiments for tonic inhibition of T-type calcium channels with the FLIPR assay was cross validated with the results from manual electrophysiology, the current ‘gold standard’ drug screening technique. Overall, the correlation between inhibitory activity in the FLIPR and patch clamp assays was excellent, with only one outlier seen in Ca_v_3.1 (AMB-CHMICA) and Ca_v_3.2 (MDMB-FUBICA) (Table 1 and Figure 1A). However, 3 of 16 SCRAs showed potent inhibition in the FLIPR assay of Ca_v_3.3 activity compared to minimal inhibition with electrophysiology. We have not established the reason for this greater discordance between the two assays of Ca_v_3.3, but it may be due to the much slower activation/inactivation kinetics of Ca_v_3.3 compared with Ca_v_3.1/3.2, or be a consequence of the protocols we used to measure Ca_v_3.3 current in patch clamp. Developing sub-type selective modulators of T-type calcium channels has proven to be very difficult due to their high sequence homology (Perez-Reyes 2003) but the FLIPR assay was able to quickly identify potent SC modulators of T-type channels and the SCRAs screened here showed some sub-type selectivity as none of the potent inhibitors of Ca_v_3.1 and Ca_v_3.2 showed any significant inhibition of Ca_v_3.3. The FLIPR assay proved to be a useful, rapid technique for preliminary screening of compounds against Ca_v_3.1 and Ca_v_3.2, the T-type calcium channels most regularly implicated in the pathogenesis human disease states.

The study which first characterized N-arachidonoylethanolamine (AEA) modulation of Ca_v_3 channels reported that the synthetic cannabinoids CP 55,940 and WIN 55,212 did not modulate Ca_v_3.1 at 10 μM, and only HU-210 (10 μM), a Δ^9^-THC analogue, had an effect (Chemin, Monteil et al. 2001). However, there are a few drugs that combine cannabinoid agonist activity and T-channel antagonist activity, these compounds produce anti-nociception in animal models of acute and neuropathic pain (You, Gadotti et al. 2011). Intriguingly, it has been reported previously that cannabinoids such as NMP-7 and cannabinoid-derived molecules such as NMP332, which block Ca_v_3.2 with IC50 values very similar to MDMB-CHMICA and AMB-CHMINACA, can produce potent anti-nociception in animal models of acute and chronic pain (Bladen, McDaniel et al. 2015). Anti-nociception was abolished when experiments were repeated in Ca_v_3.2 knockout animals, confirming the importance of the drug/channel interaction for these effects, and highlighting the potential for pain treatment with cannabinoids that also inhibit Ca_v_3.x channels. However, any future potential for SCRAs to become therapies for Ca_v_3.2 associated diseases, will likely depend on whether their agonist activity on CB1 receptors can be reduced, whilst maintaining their potency at CB2 and potent modulation of T-type calcium channels.

## Acknowledgements

This work was supported by NHMRC Project Grant 1107088 awarded to M.K., and M.C. CB was supported by a Macquarie Research Fellowship and by a Sydney Vital research grant. SM was supported by a Macquarie University Doctoral Scholarship. S.D.B. was supported by NHMRC Project Grant 1161571 and the Lambert Initiative for Cannabinoid Therapeutics, a philanthropically funded research program based at The University of Sydney.

## Author Contributions

CB designed, performed and analysed experiments and wrote the manuscript, SM performed and analysed experiments, MS created the cell lines and developed the calcium 5 assay used in FLIPR experiments, ML, MK and SB synthesised the synthetic cannabinoids, MC contributed to the design, analysis of experiments and writing of manuscript.

## ABBREVIATIONS

AEA: N-arachidonoylethanolamine
AHP: afterhyperpolarization
AMB-CHMINACA: methyl 2-((1-(cyclohexylmethyl)-1*H*-indazole-3-carbonyl)amino)-3-methylbutanoate
Ca_v_3.x: human low voltage-activated calcium channel 3.x
[Ca]_i_: intracellular calcium concentration
CB1: cannabinoid receptor type 1
CB2: cannabinoid receptor type 2
CBD: cannabidiol
Ephys: electrophysiology
FLIPR: Fluorescence Imaging Plate Reader
MDMB-CHMICA: methyl-2-[[1-(cyclohexylmethyl)indole-3-carbonyl]amino]-3,3-dimethylbutanoate
SC: synthetic cannabinoid
Δ^9^-THC: Δ^9^-tetrahydrocannabinol

